# Evolution of Gram+ *Streptococcus pyogenes* has maximized efficiency of the Sortase A cleavage site

**DOI:** 10.1101/2021.12.21.473763

**Authors:** Bradley M. Readnour, Yetunde A. Ayinuola, Brady T. Russo, Zhong Liang, Vincent A. Fischetti, Victoria A. Ploplis, Shaun W. Lee, Francis J. Castellino

## Abstract

Human plasminogen (hPg)-binding M-protein (PAM), a major virulence factor of Pattern D *Streptococcus pyogenes* (GAS), is the primary receptor responsible for binding and activating hPg. PAM is covalently bound to the cell wall (CW) through cell membrane (CM)-resident sortase A (SrtA)-catalyzed cleavage of the PAM-proximal C-terminal LPST^↓^-GEAA motif present immediately upstream of its transmembrane domain (TMD), and subsequent transpeptidation to the CW. These steps expose the N-terminus of PAM to the extracellular milieu (EM) to interact with PAM ligands, *e*.*g*., hPg. Previously, we found that inactivation of SrtA showed little reduction in functional binding of PAM to hPg, indicating that PAM retained in the cell membrane (CM) by the TMD nonetheless exposed its N-terminus to the EM. In the current study, we assessed the effects of mutating the Thr^4^ (P1) residue of the SrtA-cleavage site in PAM (Thr^355^ in PAM) to delay PAM in the CM in the presence of SrtA. Using rSrtA *in vitro*, LPS<u>Y</u>GEAA and LPS<u>W</u>GEAA peptides were shown to have low activities, while LPS<u>T</u>GEAA had the highest activity. Isolated CM fractions of AP53/ΔSrtA cells showed that LPS<u>Y</u>GEAA and LPS<u>W</u>GEAA peptides were cleaved at substantially faster rates than LPS<u>T</u>GEAA, even in CMs with an AP53/ΔSrtA/PAM[T^355^Y] double mutation, but the transpeptidation step did not occur. These results implicate another CM-resident enzyme that cleaves LPS<u>Y</u>GEAA and LPS<u>W</u>GEAA motifs, most likely LPXTGase, but cannot catalyze the transpeptidation step. We conclude that the natural P1 (Thr) of the SrtA cleavage site has evolved to dampen PAM from nonfunctional cleavage by LPXTGase.

**IMPORTANCE:** We show in this study that functional cleavage of the sortase A (SrtA) cleavage signal for M-protein, LPST*GEAA, in the Gram+ cell membrane, which allows transpeptidation of M-protein to the cell wall, as opposed to non-functional cleavage by the highly active cell membrane nonribosomal enzyme, LPXTGase, at the downstream G-residue, is highly dependent on the presence of T at position 4. From our studies, we conclude that *Streptococcus pyogenes* has evolved in a manner that maximized T at this position so that SrtA preferentially cleaved the sorting signal in order that the virulence factor, M-protein, was stabilized on the cell surface through covalent attachment to the cell wall.

## INTRODUCTION

The surface of Group A *Streptococcus pyogenes* (GAS) is decorated with a variety of proteins that function in aiding the survival and virulence of these microorganisms. Since Gram+ bacteria do not have an extracellular membrane (ECM) to stabilize surface interactions with proteins translocated from the cytosol, nor a periplasm to assist protein folding, the bacteria must employ their export channels, cytoplasmic membranes (CM), and relatively large cell walls (CW) to anchor proteins. Surface protein exposure in these cells occurs by at least three mechanisms (1): (a) moonlighting proteins bound *via* hydrophobic/charge interactions to the CW/outer capsule/techoic acids/group carbohydrate, *e*.*g*., streptococcal enolase (2, 3); (b) proteins inserted in the CM that can extend to the cell surface, *e*.*g*., lipoproteins (4); and (c) proteins covalently linked by post-translational processes to the CW, *e*.*g*., M-and M-like proteins (5). All protein epitopes exposed on the cell surface are stabilized by one or more of these processes.

Prokaryotic proteins that are ribosmally generated in the cytoplasm must be secreted or translocated into various cell compartments. This trafficking of different classes of proteins from the cytosol possesses some unique features (6). Information contained within the signal peptide and other regions of the protein sequence directs transport of proteins to their cellular compartments (7, 8). Cytosolic proteins targeted for extracellular secretion, *e*.*g*., streptokinase (SK) and the cysteine propeptidase, SpeB, are synthesized with a N-terminal cleavable signal peptide and are transported in their unfolded states through the general secretory (Sec) translocon channel, which is assembled in the CM from partner proteins. On the other hand, some surface displayed proteins, broadly classified as moonlighting proteins (9), *e*.*g*., enolase, which functions in the cytoplasm in glycolysis, contain neither a signal sequence nor other known trafficking information, but nonetheless leaves the cytosol by unknown mechanisms and is fully or partly exposed on the cell surface. These proteins are likely stabilized by hydrophobic/charge interactions with the CW/outer capsule components or can even extend from the CM to the EM.

Another group of surface displayed proteins also contain cleavable N-terminal signal sequences which terminate in a lipobox (LAAC), *e*.*g*., lipoproteins (7). These lipoproteins are enzymatically diacylated (with a diacyl glyceryl moiety) at the side-chain SH of the latent Cys^1^-residue, and then are stably inserted into the outer leaflet of the CM in Gram+ bacteria, where the signal sequence is cleaved, leaving the N-terminal Cys embedded in the CM *via* the acyl groups. This allows extension of the C-terminal regions of some of these proteins to the surface.

A final group of proteins contains a cleavable N-terminal signal sequence that directs their trafficking, and also encode a transmembrane region (TMD), near the C-terminus, *e*.*g*., M-proteins. Using this latter example, M-protein is delayed in the CM by its TMD as it emerges from the Sec channel where it is further processed by CM-embedded Sortase (Srt)-type proteases/transpeptidases, enzymes that are also located at the Sec channel (10). Housekeeping SrtA, universally present in Gram+ bacteria, recognizes a pentapeptide CW sorting signal, e.g., LPXTG, followed by a hydrophobic region of ∼21 residues and a short positively-charged cytoplasmic tail (11). Sortase A is a Cys-protease and functions by forming an acyl-enzyme complex *via* a nucleophilic attack of the active site Cys at the T^↓^ G of the CW signal followed by covalent attachment of the processed protein to the CW *via* its transpeptidase activity (12, 13). Sortases are very important to the composition and consequent properties of the CW and the cell surface, and have been used artificially to engineer the surface of cells (14). Thus, a deeper understanding of the properties of these enzymes is of great importance. It is our focus in this manuscript to explore the specificity of *Streptococcus pyogenes* SrtA-catalyzed cleavage of model peptides and, based on these results, of orthogonal M-protein variants in GAS cells that contain the functional changes. The results of this study are summarized herein.

## RESULTS

For ease in reading this manuscript, the domain structure of the Pattern D GAS-AP53 M-protein, PAM, is diagrammed in **Figure 1**. This is the only M-protein present in Pattern D GAS cells and is used for *emm* serotyping of the GAS (*emm53* in this case).

**FIG 1.**
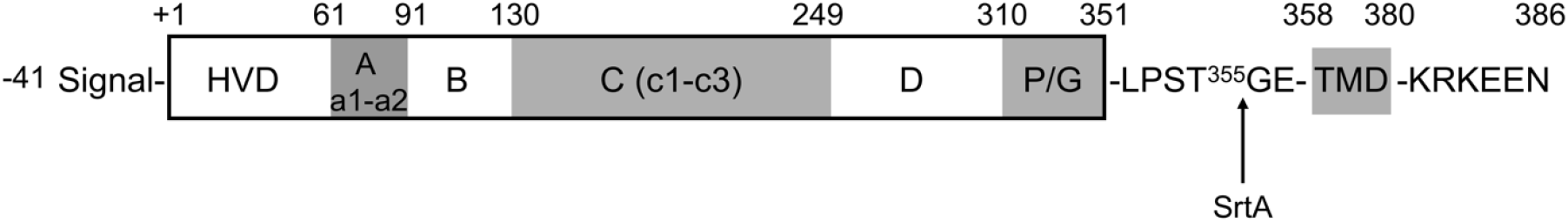
Domain structure of PAM. The 427 amino acid-residue PAM protein sequentially consisting of a 41-residue signal polypeptide, immediately downstream of which is a hypervariable region (HVR), followed by an A-domain with two hPg binding a-repeats, a B-domain, a C-domain with three c-repeats, a D domain, a Pro/Gly region, the Srt A recognition motif, a transmembrane domain (TMD) and, lastly, a short C-terminal cytoplasmic insertion sequence. Recombinant PAM spans residues 1-348. T^355^, the mutagenesis site in this study, is highlighted. PAM_short_ is a recombinant construct consisting of residues 1-132, lacking the 41-residue signal peptide), and consisting of the full HVR, the full hPg binding A-domain, and full B-domain. VEK50 is a form of PAM consisting of residues 55-104 and containing the C-terminus of the HVD, the full A-domain, and the N-terminus of the B-domain.

### Cleavage of synthetic peptides by SrtA

Recombinant (His)_6_-tagged [V^82^]SrtA and [H^37^]SrtB were expressed in *E. coli* cells, purified by affinity chromatography on columns with immobilized Ni^2+^, and eluted with imidazole. SDS gel electrophoretograms of the purified sortases are shown in **Figure 2A** and appear at the correct molecular weights. These purified enzymes were then employed to examine the cleavage by FRET of the fluorescence-tagged substrates, Dabcyl-LPS<u>X</u>GEAA-Edans, where <u>X</u> = <u>T</u>, <u>A</u>, <u>E</u>, <u>K</u>, <u>W</u>, or <u>Y</u>. The results of the assay are shown in **Figure 2B**. While all substrates are cleaved at different rates by SrtA, no linear rate of cleavage was faster than that of the native LPSTGEAA and the lowest initial rates of cleavage were found for the peptides with aromatic residues substituted at position-4 (P1), *viz*., LPS<u>Y</u>GEAA and LPS<u>W</u>GEAA. The peptides, Dabcyl-LPS<u>A</u>GEAA-Edans, Dabcyl-LPS<u>E</u>GEAA-Edans, and Dabcyl-LPS<u>K</u>GEAA-Edans displayed intermediate rates of cleavage. SrtB did not catalyze cleavage of any of these substrates.

**FIG 2.**
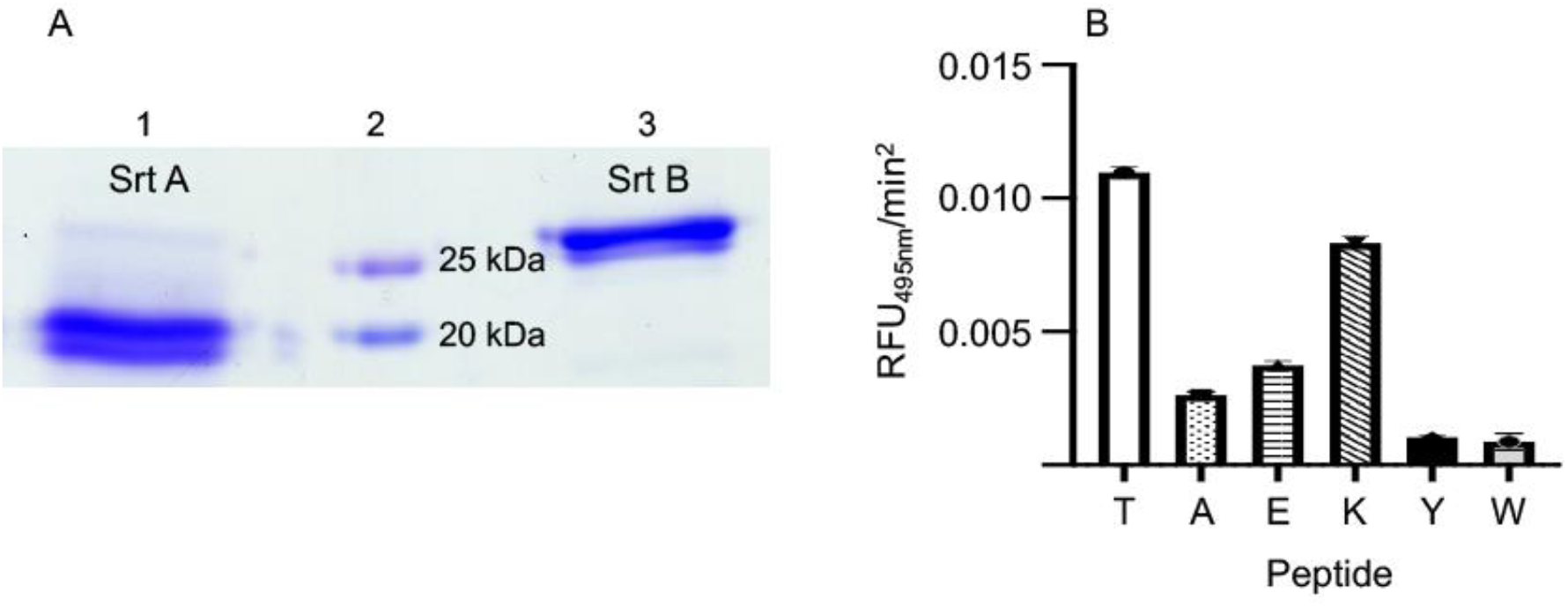
Purification and activity of recombinant sortases. **A**. SDS-PAGE gels of purified r[V^82^]SrtA and r[H^37^]SrtB. Lane 1, SrtA; Lane 2, molecular weight standards; Lane 3, SrtB. **B**. Cleavage of fluorescent peptides catalyzed by [V^82^]SrtA. Dabcyl-LPS<u>X</u>GEAA-Edans fluorescent peptides (0.2 mg), (X = T, A, E, K, Y, W) were incubated with [V^82^]SrtA and hydroxylamine and the relative fluorescence measured. Peptides (indicated on the graph by amino acid-4) were excited at 350 nm, and emission was measured at 495 nm. None of the peptides showed cleavage catalyzed by [H^37^]SrtB. N=3 for each condition.

### Subcellular distribution of PAM in isogenic AP53 cells

Since WT-PAM was selectively retained in the CM in cells in which SrtA was inactivated (15), we designed experiments to determine the cellular distribution of AP53/PAM[T^355^Y] and AP53/PAM[T^355^A], in comparison to WT-PAM, in cell lines with a targeted expression of these variants. We began by evaluating whether these SrtA-resistant PAMs were processed and secreted in the extracellular medium (EM) at mid-log phase cell growth. Using Western blot assays with our in-house polyclonal antibodies, rabbit-anti-PAM and rabbit-anti PAM_short_, we did not find meaningful differences in the levels secreted into EM between WT-PAM (lane 2) and the PAM variants (lanes 3 and 4) of these different cell lines **(Figure 3A)**. Moreover, since the Western blot analyses were performed using anti-PAM_short_, the blot clearly shows that the secreted PAM variants contain the N-terminus of PAM. The molecular weights of the PAM variants are the same as that of WT-PAM, and all are nearly identical to recombinant PAM (lane 6), with no band appearing for the AP53/*Δpam* cells (lane 5), as anticipated. These results demonstrate that PAM[T^355^Y] and PAM[T^355^A] have been cleaved by an enzyme that recognizes the SrtA cleavage sequences, LPS<u>A</u>GEAA and LPS<u>Y</u>GEAA.

**FIG 3.**
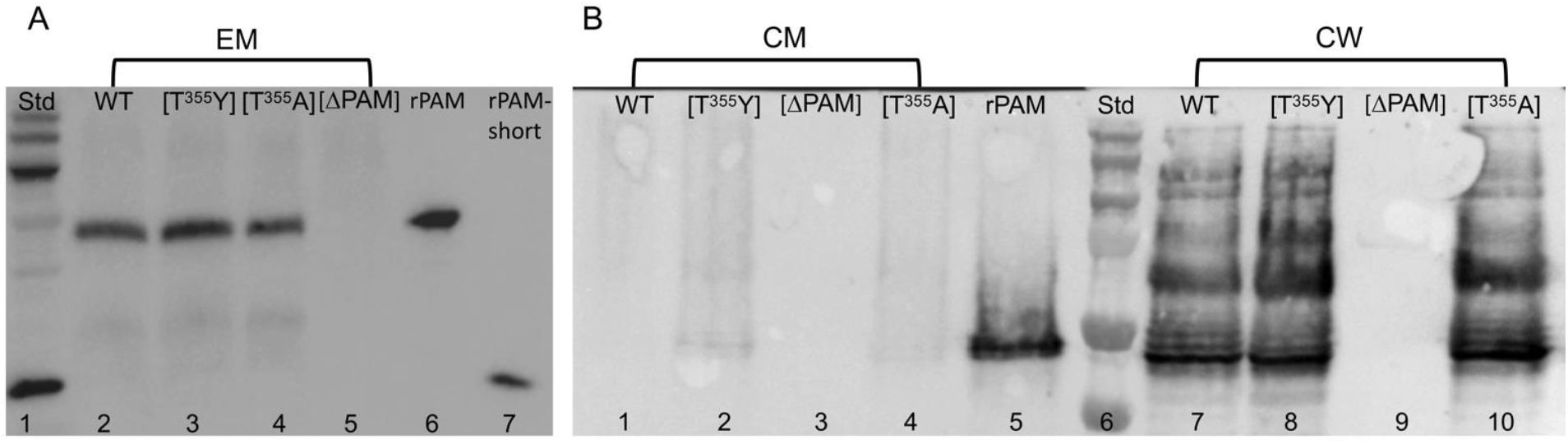
PAM distributions in fractionated mid-log phase cells. **A**. Western blot analysis of the extracellular medium (EM) with anti-PAM_short_. The EM samples were concentrated 40x prior to analysis. Lane 1, Mol. Wt. ladder; Lanes 2-5 contained EMs (40x concentrated) from: Lane 2, WT-AP53; Lane 3, AP53/PAM[T^355^Y]; Lane 4, AP53/PAM[T^355^A]; Lane 5, AP53/ΔPAM. Lane 6 is rPAM and Lane 7 is PAM_short_. **B**. AP53 cells were washed and treated with 20 µg Ply-C for 1 hr. Cell wall (CW) and cytoplasmic membrane (CM) fractions were analyzed by Western blotting with anti-PAM_short_. Lanes 1-4 represent the CM fractions from: Lane 1, WT-AP53; Lane 2, AP53/PAM[T^355^Y]; Lane 3, AP53/ΔPAM; Lane 4, AP53/PAM[T^355^A] cells. Lane 5 is rPAM, and Lane 6 is a standard molecular weight ladder. Lanes 7-10 are CW fractions from: Lane 7, WT-AP53; Lane 8, AP53/PAM[T^355^Y]; Lane 9, AP53/ΔPAM; Lane 10, AP53/PAM[T^355^A] cells.

We next examined the distribution of PAM in mid-log phase subcellular fractions of WT-AP53, AP53/PAM[T^355^A], and AP53/PAM[T^355^Y]. The results of the experiment are shown in **Figure 3B**. As expected, PAM was not present in AP53/ΔPAM (lane 3) cells, and there is very little PAM in the CM fractions obtained from WT-AP53 (lane 1), AP53/PAM[T^355^Y] (lane 2), and AP53/PAM[T^355^A] cells (lane 4). However, PAMs were demonstrated in the CW fractions of all isogenic strains, except for AP53/ΔPAM (lanes 7-10). Thus, the PAM variants were fully cleaved, despite the reduced level of activity that the synthetic peptides showed against SrtA *in vitro*. Further, these peptides were covalently attached to the CW, as indicated by the higher molecular weight PAM+ bands that showed a distribution of PAM with various amounts of partially Ply-C-digested CW peptidoglycan, similar to WT cells. This demonstrates that the PAM variants are processed through SrtA. To explain these results, it has been suggested that SrtA *in vitro* has a lower level of activity toward its cleavage site in synthetic peptides than SrtA in the CM since neither the peptides, nor SrtA, are imbedded in the CM in the *in vitro* assays (16). While the SrtA cleavage site peptide variants may have reduced activity toward SrtA *in vitro*, the data show that the SrtA activity is sufficient in cells to both cleave and transpeptidate these variants on the CW when these sequences are present in PAM.

### Expression of wt-PAM and its Srt motif variants on the surface of isogenic AP53 cells

To confirm the presence and integrity of the PAM variants on the AP53 cell surface, we performed whole cell immunoassays using anti-PAM_short_, as well as FCA analysis of hPg binding to the cell lines. The data of **Figure 4** show that equivalent amounts of each PAM are present on AP53 cells with their N-termini largely intact, as revealed by their reactivities toward rabbit-anti-PAM_short_ when whole cells are analyzed by FCA (**Figure 4A**) and by ELISA **(Figure 4B)**. Further, the cell surface PAM fully reacts with hPg (**Figure 4C**), a property of the N-terminal A-domain. We also show that each of the PAM variants stimulate the activation of hPg by SK2b **(Figure 4D,E)**, a SK subform that only enhances hPg activation when hPg is bound to PAM, and which requires the N-terminal A-domain of PAM to be exposed to the extracellular solution (17, 18). These results demonstrate that PAM and its variants interact in an equivalent and functional manner with hPg.

**FIG 4.**
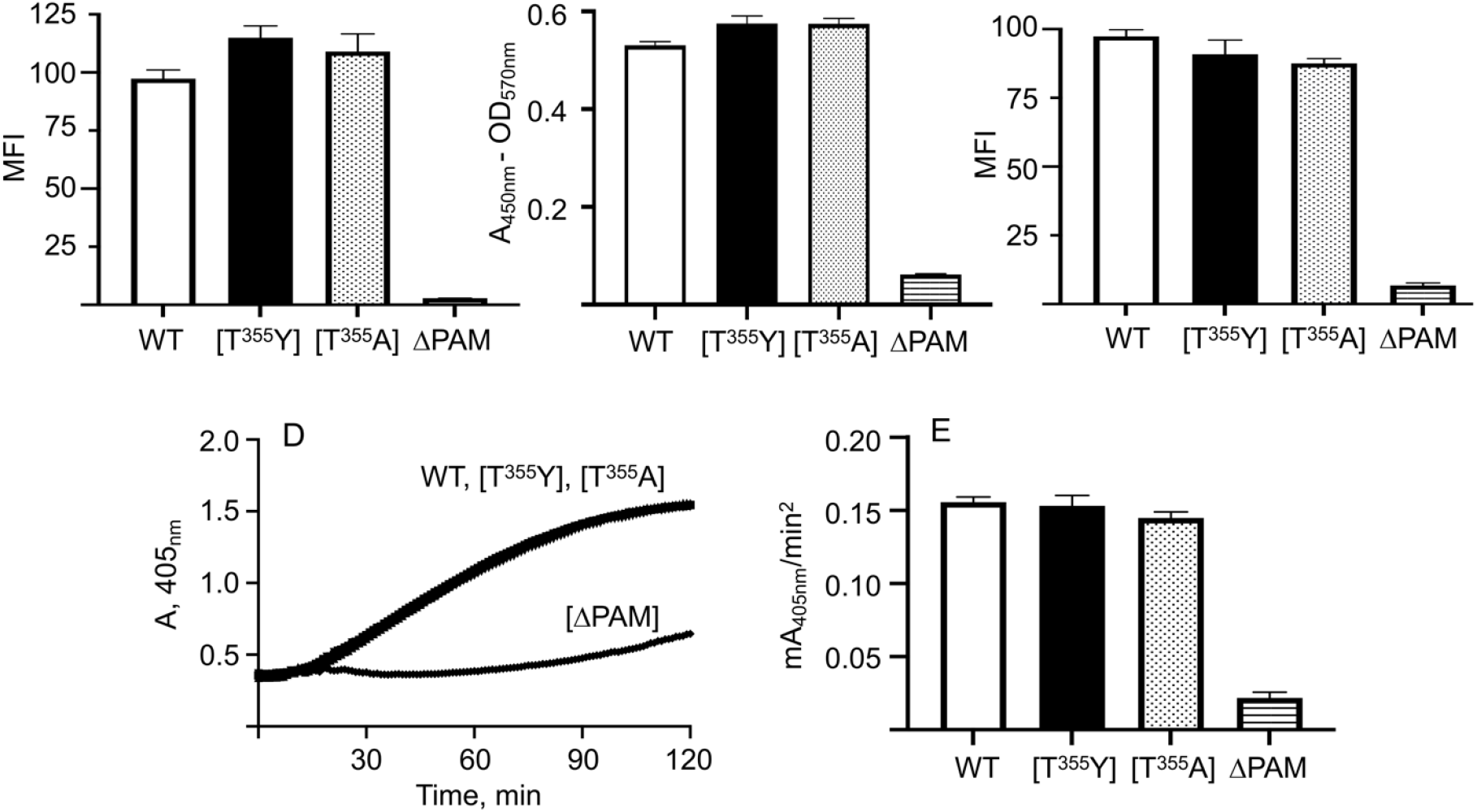
Binding and activation of hPg by variant PAMs. **A**. *FCA of rabbit-anti-PAM binding to whole AP53 cells*. WT-AP53, AP53/PAM[T^355^Y], AP53/PAM[T^355^A], and AP53/ΔPAM mid-log-phase cells were incubated with rabbit-anti-PAM_short_ followed by Alex Fluor 488-chicken-anti-rabbit IgG. The median fluorescence intensity (MFI) determined by FCA is shown as bar graphs of the relative levels of PAM on each of the different cell lines. **B**. *ELISA of rabbit-anti PAM*_*short*_ *binding to whole AP53 cells*. Incubation with rabbit-anti PAM_short_ as in **A**, followed by goat-anti-rabbit IgG-HRP. Equal numbers of cells were added to wells of a 96-well plate, and the HRP substrates, H_2_O_2_ and TMB were added to each well. The reaction was terminated with H_2_SO_4_. The OD_570nm_ was determined and subtracted from A_405nm_. **C**. *hPg binding to isogenic AP53 strains*. Mid-log phase AP53 cells were treated with hPg and incubated with monoclonal mouse-anti-hPg IgG. The bound hPg was detected by incubation with Alex Fluor 488-donkey-anti-mouse IgG and analyzed as in **A. D**. *hPg activation by SK2b*. The isogenic AP53 cells were placed in individual wells of a BSA-blocked 96-well microtiter plate, after which hPg was added, followed by S2251 and, lastly, SK2b. The hPm-catalyzed release of p-nitroaniline from S2251 was continuously monitored at 405 nm for 120 min. **E**. *hPg activation by SK2b*. The initial rates of activation calculated from these curves are represented as bar graphs. At least three different biological replicates were used for each strain. The buffer was 10 mM Na-Hepes/150 mM NaCl, pH 7.4, at room temperature. N ≥ 3 for each experiment.

### Cleavage of peptides by subcellular fractions from SrtA-depleted AP53 cells

We expanded this study to ascertain the subcellular fraction responsible for the cleavage of PAM[T^355^Y]. Here, we used three additional isogenic AP53 GAS strains: 1) AP53/ΔSrtA; 2) a double mutant deficient in both SrtA and SrtB, (AP53/*ΔsrtA*/*ΔsrtB*); and 3) a mutant deficient in SrtA and carrying the SrtA cleavage site mutation in PAM (AP53/*ΔsrtA*/PAM[T^355^Y]). We first determined the subcellular fraction of the WT-AP53 cells that catalyzed cleavage of the WT-sequence, Dabcyl-LPS<u>T</u>GEAA-Edans, and its variant, Dabcyl-LPS<u>Y</u>GEAA-Edans. From the results of **Figure 5A**, at dilutions wherein the native peptide is cleaved to a low extent by all of the cell fractions, the same concentration of variant peptide is very rapidly cleaved by a component(s) in the CM. Interestingly, in CMs from cells in which both SrtA and SrtB are inactivated (AP53/ΔSrtA/ΔSrtB), the variant peptide is rapidly cleaved under conditions wherein the peptide with the WT-sequence is cleaved slowly (**Figure 5B**). Thus, the responsible protease(s) has a much higher specificity for aromatic residues at the 4^th^ (P1) position of the SrtA cleavage site than the Thr at this same position in of the WT-peptide.

**FIG 5.**
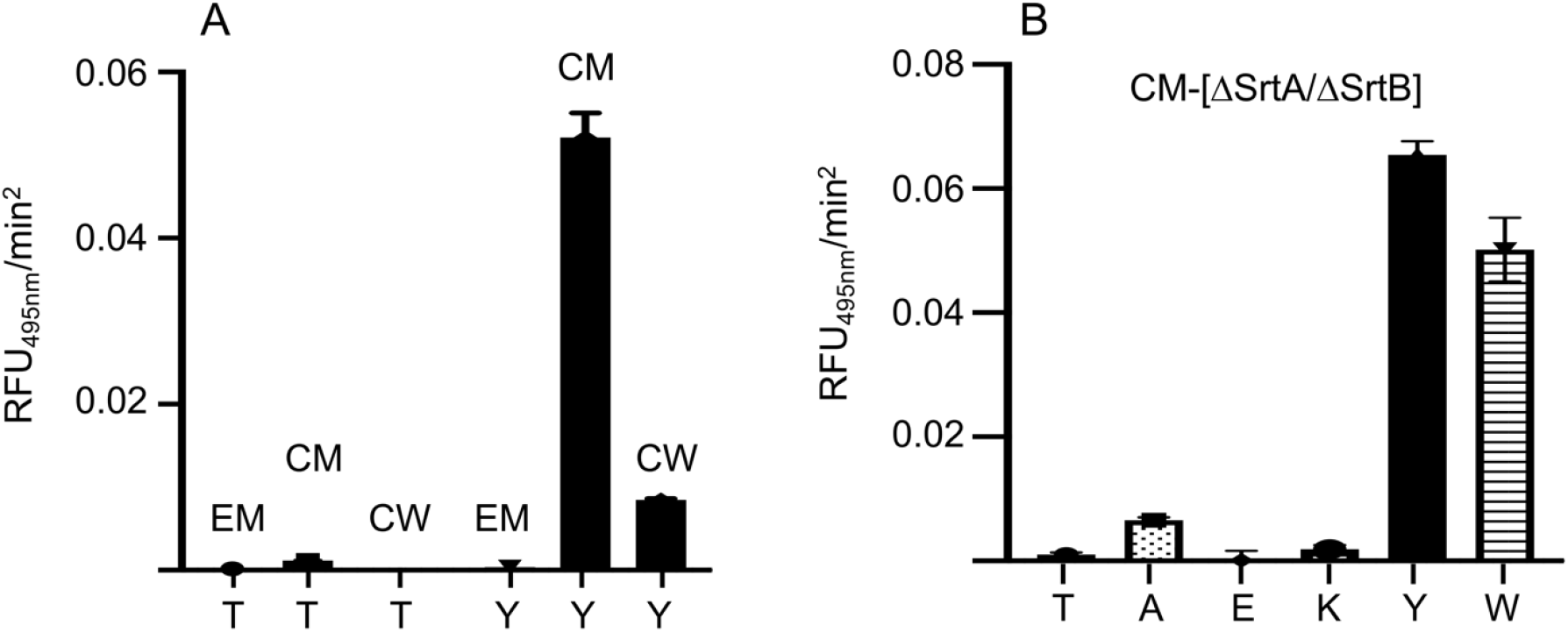
Cleavage of synthetic peptides with AP53 subcellular fractions. **A**. Cleavage of the synthetic peptides, Dabcyl-LPS<u>T</u>GEAA-Edans (T) and Dabcyl-LPS<u>Y</u>GEAA-Edans (Y) with cell supernates (EM), cell membranes (CM) and cell wall (CW) fractions of WT-AP53 cells. **B**. Synthetic peptide cleavage by AP53/ΔSrtA/ΔSrtB CM fractions. Dabcyl-LPS<u>T</u>GEAA-Edans (T), Dabcyl-LPS<u>A</u>GEAA-Edans (A), Dabcyl-LPS<u>E</u>GEAA-Edans (E), Dabcyl-LPS<u>K</u>GEAA-Edans (K), Dabcyl-LPSYGEAA-Edans (Y), and Dabcyl-LPSWGEAA-Edans (W) peptides (0.2 mg) were dissolved in 50 mM Tris/50 mM NaCl, pH 7.4. The peptide solution was added to AP53/ΔSrtA/ΔSrtB-derived membrane fractions and incubated for 3 hr with the CM fraction of these cells. In all cases, the peptides were excited at 350 nm and emission was measured at 495 nm. N=3 for each condition.

To further investigate the cleavage mechanism, we assessed PAM distribution in subcellular fractions of AP53/*ΔsrtA* and AP53/*ΔsrtA*/*pam[T*^*355*^*Y]*. The data of **Figure 6A** demonstrate that AP53/ΔSrtA/Pam[T^355^Y] cells secrete into the EM high levels of PAM[T^355^Y] that has been cleaved in a manner that displayed a very similar molecular weight as rPAM and WT-PAM secreted by WT-PAM cells. Whereas PAM[T^355^Y] is cleaved from the CM (**Figure 6B**), as evidenced its molecular weight, it is not covalently linked to the CW, since partially-digested CW bands containing PAM[T^355^Y] are not observed. Further, PAM[T^355^Y] is retained in the CM in the absence of SrtA, whereas WT-PAM is not (**Figure 6C**). Overall, these experiments suggest that while SrtA is not required for cleavage near the SrtA site of PAM, SrtA is necessary for transpeptidation to occur.

**FIG 6.**
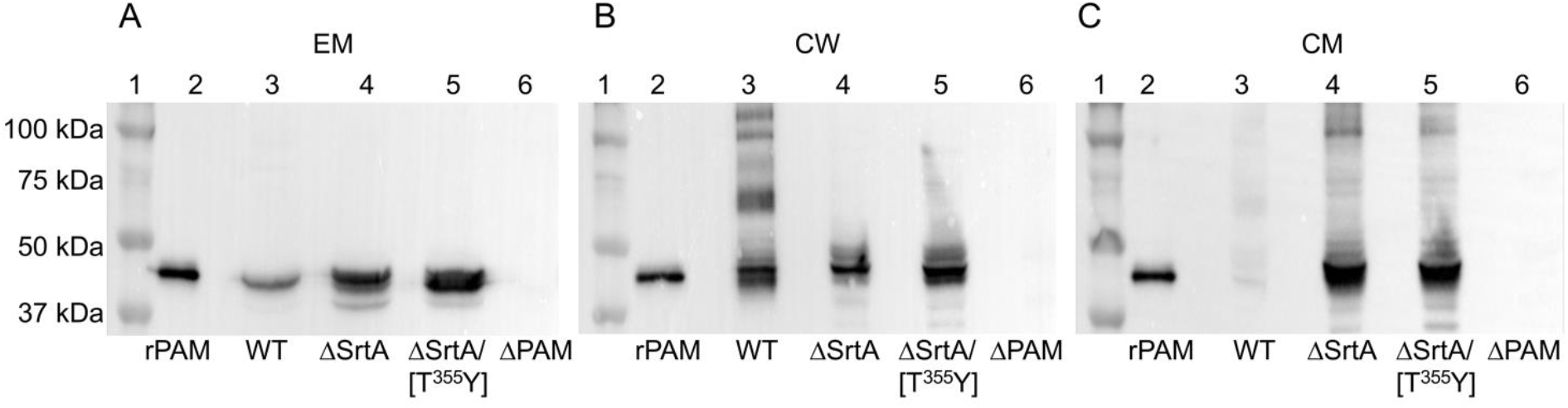
Western blot analysis for PAM in AP53 cell fractions. Mid-log phase isogenic AP53 strains were grown to mid-log phase and (**A**) EM - Cell supernate (40X concentrated), (**B**) CW - cell wall, and (**C**) CM - cell membrane separated were separated using 10% tris-glycine SDS gels, transferred to PVDF membranes, visualized with anti-PAM_short_. Lane 1, molecular weight standards; Lane 2, rPAM; Lane 3, WT-AP53; Lane 4, AP53/ΔSrtA; Lane 5, AP53/ΔSrtA*/*PAM[T^355^Y]; Lane 6, AP53/ΔPAM.

### The variant PAM[T^355^Y] is functionally displayed on the AP53 cell surface

We further show that PAM from AP53/ΔSrtA and AP53/ΔSrtA/PAM[T^355^Y] cells is displayed on the cell surface in the absence of SrtA. This was assessed by examining their interaction with anti-PAM_short_, which also indicated that all PAMs possessed a relatively intact N-terminus (**Figure 7A**). The surface exposed PAMs were also functional in stimulating hPg activation by SK2b (**Figure 7B**). Whether the PAM extending from the CW and/or from the CM is the source of the surface-exposed PAM is not known at this stage, but a SrtA deficiency does not significantly affect the ability of PAM to function in binding and stimulating the activation of hPg in whole AP53 cells (15).

**FIG 7.**
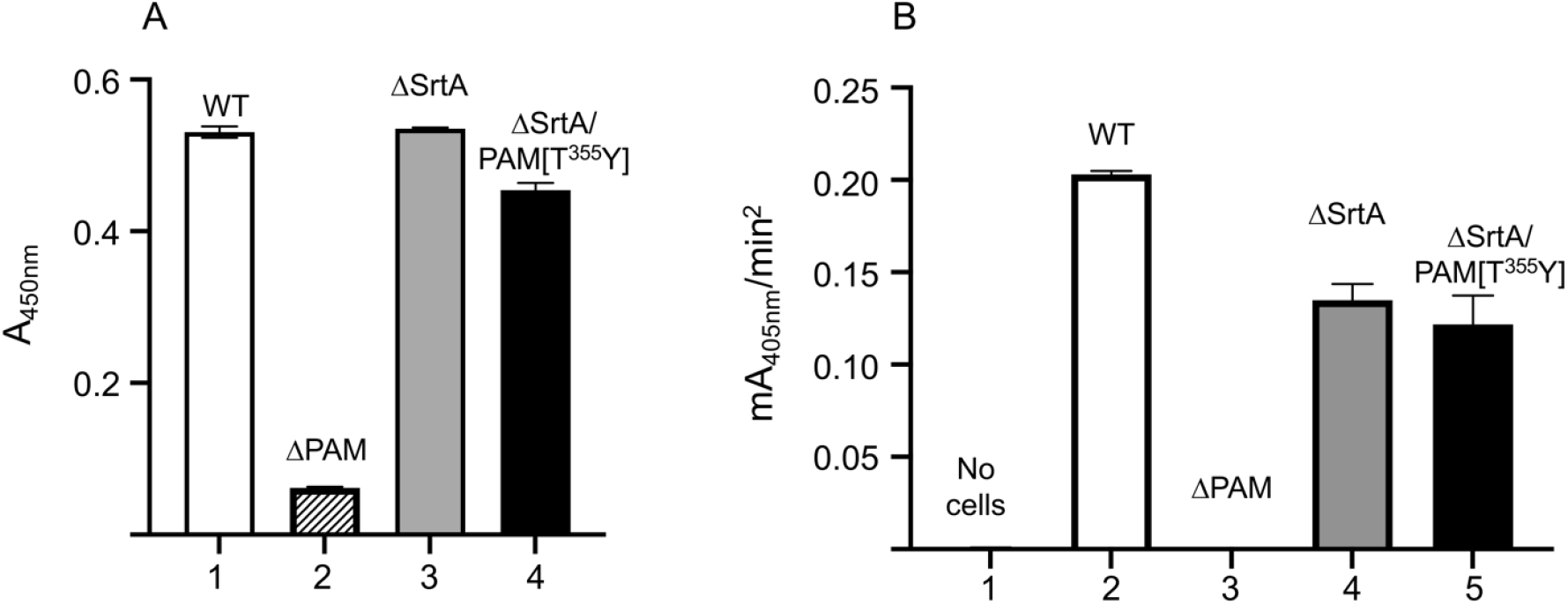
Functional presence of PAM on whole cells. **A**. Binding of anti-PAM to whole cells by ELISA. Mid-log phase isogenic AP53 cells were incubated with rabbit-anti PAM_short_ and goat-anti-rabbit IgG-HRP and developed with H_2_O_2_ and TMB. The reaction was terminated with H_2_SO_4_ and the OD_570nm_ (light scatter) was subtracted from the A_450nm_. Lane 1, WT-AP53; Lane 2, AP53/ΔPAM; Lane 3, AP53/ΔSrtA; Lane 4, AP53/ΔSrtA/PAM[T^355^Y]. **B**. Stimulation of SK2b-catalyzed hPg activation by mid-log isogenic AP53 strains. Isogenic AP53 cells were placed in individual wells of a BSA-blocked 96-well microtiter plate, after which hPg was added, followed by S2251/SK2b. The hydrolysis of S2251 was continuously monitored at A_405nm_. At least three different clonal cells were used for each strain. The buffer was 10 mM Na-Hepes/150 mM NaCl, pH 7.4, at room temperature. The initial rates of activation are presented as bar graphs: Lane 1, no cells; Lane 2, WT-AP53; Lane 3, AP53/ΔPAM; Lane 4, AP53/ΔSrtA; Lane 5, AP53/ΔSrtA/PAM[T^355^Y] cells.

The SEM results (**Figure 8**) confirm all the findings of this manuscript. The micrographs **(8A, B)** are negative controls showing that PAM is not detected on AP53 cells when anti-PAM is not present (**Figure 8A**) or in cells with anti-PAM but with a targeted deletion of the *pam* gene (**Figure 8B**). PAM is visible with anti-PAM_short_ in WT-AP53 (**Figure 8C**) cells and in AP53/PAM[T^355^Y] cells (**Figure 8D**). Finally, and importantly, WT-PAM (**Figure 8E**) and PAM[T^355^Y] (**Figure 8F**) are abundant on the cell surface when the gene for *srtA* is genetically inactivated.

**Figure 8.**
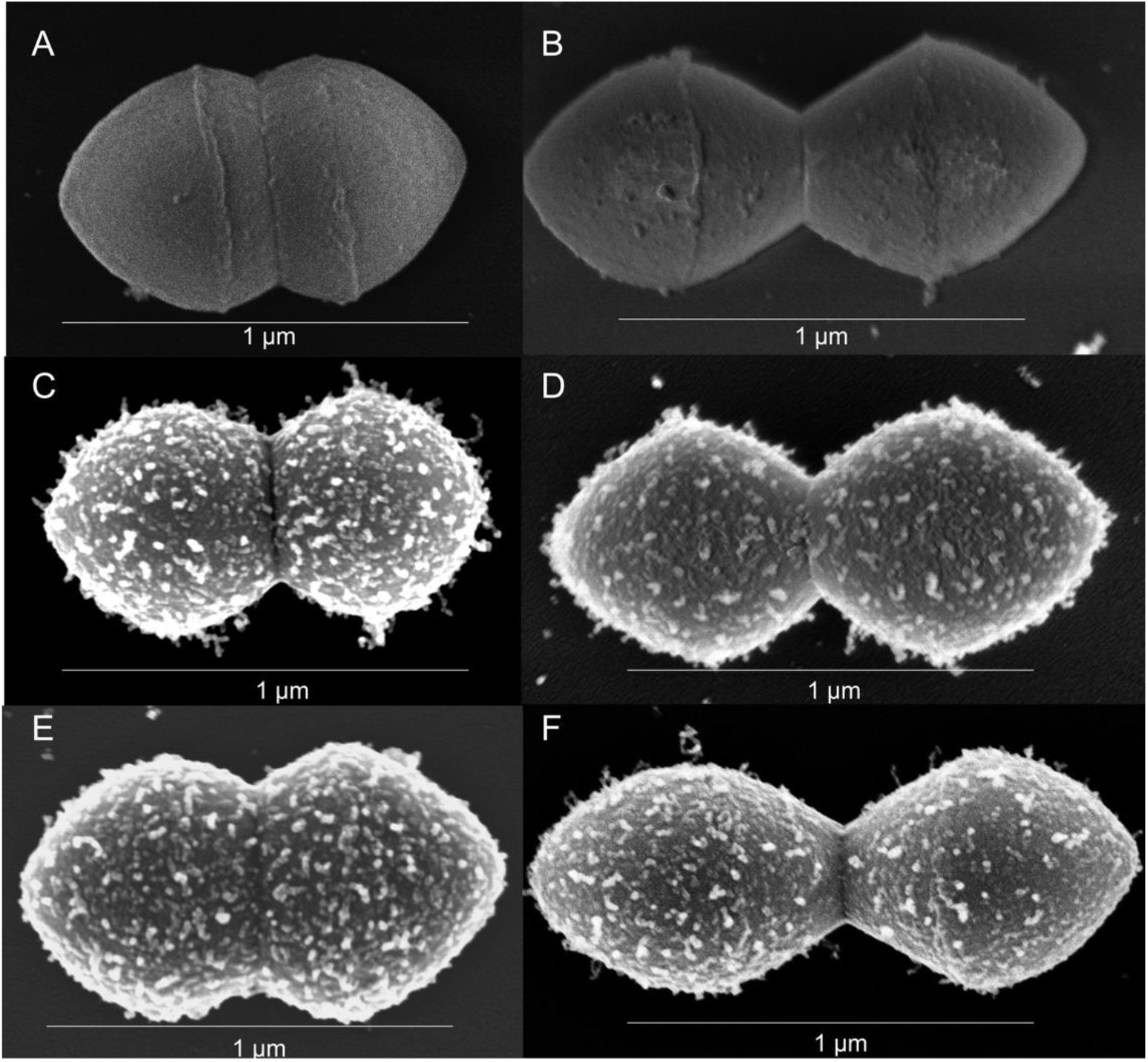
SEM images of AP53 GAS cells grown to mid-log phase and incubated with rabbit-anti PAM_short_ to determine proximity and relative abundance of PAM on the surface of isogenic AP53 strains. **A**. WT-AP53, no antibody; **B**. AP53/ΔPAM with rabbit-anti PAM_short_; **C**. WT-AP53 with rabbit-anti-PAM_short_; **D**. AP53*/*PAM[T^355^Y] with rabbit-anti PAM_short_; **E**. AP53/ΔSrtA*/*PAM[T^355^Y] with rabbit-anti PAM_short_; **F**. AP53/ΔSrtA with rabbit-anti PAM_short_.

## DISCUSSION

Since Gram+ bacterial strains do not have an outer membrane to stabilize surface-exposed proteins, specialized mechanisms of protein trafficking exist such that surface proteins are expressed in functional forms on the cell surface. The relatively extensive CW in these bacterial strains is of great utility in this regard. The study of surface proteins of bacterial pathogens is critical to their potential utility as vaccine candidates (19). In particular, vaccine candidates toward GAS have largely involved M-protein variants and other surface-exposed proteins as potential antigenic determinants for robust antibody-mediated responses (20-22). Considering our results, it is clear that further mechanistic studies are needed to better understand how GAS surface proteins, especially M-protein and PAM-protein variants, can be differentially localized and exposed to outer surfaces and how variant LPXTG motifs may be a major mechanism by which GAS can quickly vary outer surface localization to increase antigenic variation.

To examine the surface proteins in GAS isolate AP53, we mined the genomic sequence of this strain (23). A total of 28 candidate lipoproteins with a defined lipobox were found. The lipoproteins are imbedded in the CM by post-translational N-terminal anchoring, that also enable display of surface proteins by extension of their C-terminal regions through the CW and outer capsule to the EM (24). Additionally, several moonlighting proteins, especially those involved in glycolysis, *e*.*g*., enolase and GAPDH, are cytosolic proteins without signal sequences, but are also found in the CW and on the AP53 cell surface (2, 3, 25). Their trafficking pathways from the cytosol are largely unknown. We also identified ∼17 proteins that contain the SrtA signal sequence, one with an Ala at P1 of the SrtA cleavage site and the remaining 16 proteins possessing Thr at P1 of this cleavage site. Biologically relevant cleavages of the Tyr^4^ or Trp^4^ mutants at or near the P1 site was heretofore unknown.

We show that cells containing the PAM[T^355^Y] mutation display the same mid-log phase subcellular distribution as cells containing WT-PAM, and this variant is covalently attached to the CW. The EM also contains both WT-PAM and PAM[T^355^Y] at molecular weights that suggest that these proteins have been similarly cleaved. We presume that while Tyr^4^ at the P1 site is poorly cleaved by SrtA *in vitro*, cleavage by SrtA in cells occurs at a sufficient rate to place this mutant on the CW. Importantly, the SrtA-bound PAM[T^355^Y] is transpeptidated to approximately the same extent as the natural PAM[T^355^].

The CM spanning Sec translocon, consisting of a SecYEG protein triheteromer, as well as ancillary proteins, including SecA, transports unfolded proteins, *e*.*g*., PAM, from the cytoplasm through the CM, where other processing proteins, including signal peptidases and SrtA, can function to deliver the protein to its final destination (26). We show that when SrtA is inactivated, PAM is delayed in the CM where it is nonetheless cleaved by another specific protease, within the SrtA site, but is not transpeptidated to the CW. In this case, PAM is found noncovalently bound within the CM and also the EM where it is functional regarding hPg binding and activation. We hypothesized that PAM delayed in the CM nonetheless spans the CW and presents its N-terminus to the EM, where hPg interacts with the exposed A-domain and is activated by SK2b secreted by the cells (15). Also, the PAM cleaved from the CM and present in the CW is likely noncovalently bound to peptidoglycan and can also expose its A-domain for the same purpose.

Since a SrtA inactivation affects many other proteins of GAS, we sought another manner of inhibiting cleavage of the SrtA site specifically in PAM that would support our hypothesis. PAM is cleaved between the P1 (Thr^4^) and P1’ (Gly^5^) position (at ^↓^) of the LPS<u>T</u>^↓^GEAA consensus sequence of PAM, immediately upstream of its TMD near the C-terminus. Thus, we sought a peptide sequence that would be highly attenuated toward SrtA cleavage, specifically in PAM. We first investigated the activation rates catalyzed by SrtA of a series of LPS<u>X</u>GEAA peptides *in vitro* and found that LPS<u>T</u>GEAA is the most rapidly cleaved peptide, as expected. The next best peptide sequence contains Ala at P1 of the SrtA consensus cleavage site, a known biologically relevant residue in PAM SrtA-sensitive proteins. Other peptides are cleaved at slower rates, with LPS<u>Y</u>GEAA and LPS<u>W</u>GEAA as the most unfavorable substrates for rSrtA. Thus, we generated an AP53 cell line with a targeted replacement of WT-PAM by PAM[T^355^Y]. Surprisingly, we found that the variant PAM was distributed in AP53 cellular fractions similarly to WT-PAM, in that PAM[T^355^Y] was transpeptidated to the peptidoglycan to approximately the same extent as WT-PAM. This shows that the low level of activity of the LPS<u>Y</u>GEAA peptide toward SrtA *in vitro* was nonetheless sufficient *in vivo* for the SrtA reactions to occur on the CM.

A different picture emerges when isolated CMs from WT-AP53 cells are compared to AP53/ΔSrtA/ΔSrtB cells in their ability to cleave the synthetic peptides, LPS<u>T</u>GEAA, LPS<u>Y</u>GEAA, and LPS<u>W</u>GEAA. Here, the CM cleaves within the SrtA site, but does not transpeptidate the PAMs to the CW. An explanation for these finding lies with the unique membrane endopeptidase, LPSTGase, which cleaves within the SrtA site at P2 (Ser^3^) and P2’ (Glu^6^) of the WT-sequence, (Ser^354^ and Glu^357^ of PAM) (27). Whether this enzyme participates in transpeptidation is not known but, according to our findings, this is unlikely to be the case.

LPXTGase is an unusual enzyme in that it is very hydrophobic, heavily glycosylated, and contains unusual unidentified amino acids as well as D-alanine of the CW. It is believed that this enzyme is nonribosomally produced and is generated during assembly of the CW (27). According to our results herein, we conclude that this enzyme prefers aromatic residues at the 4^th^ position of a SrtA cleavage sequence and has low activity when the natural Thr is present at this location. Thus, it is tempting to conclude that GAS has evolved to greatly prefer Thr at this location in order that the SrtA can preferentially function to properly process proteins that emerge through the Sec channel, and avoid nonfunctional cleavage by LPXTGase.

The conclusions reached in this study are verified by the SEM images of the isogenic cell lines. WT-AP53 cells react with anti-PAM_short_, showing an abundance of PAM on the cell surface, as is also the case with AP53/ΔSrtA. This confirms that through its elongated structure, the PAM N-terminal hPg-binding region reaches the cell surface, while its C-terminus is trapped both in the CM and cross-linked peptidoglycan. The SEM scan similarly shows that PAM[T^355^Y] also extends through and to the CW, exposing its N-terminus to the extracellular medium with or without SrtA. Thus, while SrtA is necessary to permanently link PAM to the peptidoglycan (28), this step is not required to expose PAM on the cell surface to bind and activate hPg, though this latter exposure may be transient. We speculate that LPXTGase, a membrane-bound protein like SrtA (29), may be a primordial nonribosomally-produced enzyme that has evolved with Gram+ bacteria for critical, yet unknown functions. Its location and specific endopeptidase activity would have a negative effect on bacterial survival through nonproductive cleavage within the anchor region of M- and M-like proteins. To counter this, we reason that the bacteria ultimately evolved a Thr (or Ala) at the SrtA LPXT^G cleavage site, thereby reducing the negative effects of LPXTGase.

## MATERIALS AND METHODS

### Isogenic AP53 strains

GAS isolate AP53 (*emm53*) was obtained from Dr. G. Lindahl, Lund, SE and used as the parent strain.

The generation of the isogenic AP53 strain with targeted deletion of the *srtA* gene, *via* allelic replacement with the *cat* gene, has been described earlier (15, 30).

Allelic replacement of the 726 bp *srtB* gene with the *cat* gene was similarly conducted. The targeting plasmid contained the *cat* gene (660 bp), flanked by 468 bp upstream of the ATG signal for *srtB* and 378 bp downstream of its TAA stop codon. During this construction, 5’-*NotI* and 3’-*XhoI* restriction sites were also cloned into this DNA segment using PCR primers and employed for insertion into the same sites of the temperature-sensitive plasmid pHY304 (from M. J. Walker, Queensland, AU), which also contained the downstream erythromycin resistance (*erm*) gene. Integration of *cat* into the *srtB* chromosomal locus was accomplished by single crossover (SCO) at 30° C and double crossover (DCO) at 37° C, which also eliminated the *erm* gene. Screening of confirmed SCO colonies was conducted by erythromycin resistance. DCO colonies obtained from the confirmed SCO colonies were screened by loss of erythromycin resistance after loss of the *erm* gene resulting from the DCO. DCO colonies were further tested by resistance to chloramphenicol. The substitution of *cat* for *srtB* was further confirmed by PCR of the genomic DNA with *cat* and *srtB* specific primers.

The AP53/*ΔsrtA/ΔsrtB* double deletions were constructed as above, with the targeting plasmid containing *ΔsrtB* incorporated into the AP53/*ΔsrtA* cells by DCO as above.

The same general procedures were used for generation of AP53/*pam[T*^*355*^*A]*, AP53/*pam[T*^*355*^*A]*, and AP53/*pam[T*^*355*^*Y]* in AP53 cells or in AP53/*ΔsrtA* cells. The DCO colonies were screened using primers specific for the LPSAG and LPSYG motifs. Colonies were analyzed by Sanger sequencing for final confirmation.

### Proteins

#### Cloning and expression of recombinant SrtA and SrtB

SrtA (GenBank: AMY97447.1), residues 82-249 ([V^82^]SrtA), without its predicted N-terminal signal peptide nor its TMD and peripheral membrane region, was cloned from GAS-AP53 genomic DNA (GenBank: CP013672.1) using a 21 bp forward primer beginning at the first nucleotide encoding V^82^ and a 30 bp reverse primer beginning at the stop codon for *srtA*. The reverse primer included introduction of a new *BamH1* site after the *srtA* stop codon. PCR reactions were carried out with Phusion Hot Start High-Fidelity DNA Polymerase (New England Biolabs, Ipswich, MA). After digestion with *BamH1*, the PCR products were ligated into *PshA1/BamH1*-digested pET42a (EMD Biosciences, Darmstadt, Germany), yielding the *srtA* cDNA with an additional N-terminal 276-amino acids from pET42a. The entire expression cassette contained, sequentially, a glutathione sulfur-transferase (GST) tag - a 15 residue S-tag (31) - a (His)_6_ tag for purification - a Factor Xa (FXa) cleavage site (IEGR^↓^) - followed by [V^82^]SrtA.

The same procedure was used to prepare the plasmid for *srtB* expression (GenBank: AMY96673.1), lacking the N-terminal 36-residue signal peptide and its TMD, and providing residues 37-241 ([H^37^]SrtB).

The plasmids were expressed in *Escherichia coli* BL21/DE3 cells as we have described previously for proteins such as PAM, PAM fragments, and SK (18, 32, 33). After purification by Ni^+^-charged His-trap resin (EMD Biosciences), elution with imidazole, and cleavage of the FXa-sensitive site (at R), the final purified SrtA proteins consisted of [V^82^]SrtA or [H^37^]SrtB, without non-native amino acids (18).

#### Human plasminogen (hPg) binding Group A streptococcal M-protein (PAM)

PAM was expressed in *Escherichia coli* and purified as described. The final PAM product consisted of residues 1-358 (**Figure 1)**.

#### Streptokinase cluster 2b (SK2b)

The SK2b variant which is coinherited with PAM was also expressed in *E. coli* as previously described (18).

#### Human plasminogen (hPg)

hPg was expressed in *Drosophila Schneider S2* cells and purified by Sepharose-lysine affinity chromatography (34).

#### Antibodies

Polyclonal antibodies were generated against PAM in rabbits by standard procedures. Similarly, a rabbit polyclonal antibody was generated against a form of PAM containing only the intact N-terminal 132 amino acids (PAM_short_; residues 1-132, lacking the 41-residue signal peptide). This PAM fragment contained the HVR +1, the full hPg binding A-domain, and the B-domain. Anti-PAM_short_ showed no reactivity with VEK50, a form of PAM (residues 55-104) that contains the C-terminus of the HVD, the full hPg binding A-domain, and the N-terminus of the B-domain. This implies that anti-PAM_short_ requires the HVR for reactivity.

### Genotyping

Genomic DNAs (gDNA) were prepared from single colonies of AP53 lines selected from sheep blood agar plates. The colonies were treated with lysozyme/proteinase K and suspended in 100 mM Tris/5 mM EDTA/0.2% SDS/200 mM NaCl, pH 8.5. gDNAs were extracted with phenol/chloroform/isoamyl alcohol (25/24/1), precipitated with isopropanol, and washed with 70% ethanol in nuclease-free H_2_O.

PCR was employed with gDNA from the various strains to determine whether the desired gene alterations were present in the genomes. To detect the *srtA* gene, 24 bp internal primers that consisted of a forward primer *(srtA-F)* beginning at nucleotide 233 of *srtA* and a reverse primer *(srtA-R)* beginning at nucleotide 492 of *srtA*, were used. The *srtA* gene showed a 260 bp amplicon with these primers. For detection of the *cat* gene, 20 bp and 21 bp primers were employed beginning at nucleotide 241 for the forward primer (*cat-F*) of the *cat* DNA and at nucleotide 460 for the reverse primer (*cat-R*) began of the *cat* cDNA. This provided a 220 bp amplicon when the *cat* gene was present.

To detect the *srtB* gene, a 19 bp forward primer (*srtB-F*) beginning at nucleotide 82 and a 20 bp reverse primer (*srtB-R*) beginning at nucleotide 317 was used to yield an amplicon of 236 bp.

For detection of *WT*-*pam*, an 18 bp forward primer (*pam-F*), beginning at nucleotide 81 of the *pam* gene and a 20 bp reverse primer (*pam-R*) which began at 439 bp of the *pam* gene yielded a 359 bp amplicon when *pam* was present. For detection of *pam[T*^*355*^*A]*, a 22 bp forward primer (*pam[T*^*355*^*A]-F)* beginning at nucleotide 1081, and a 23 bp reverse primer (*pam[T*^*355*^*A]-R)* beginning at nucleotide 1387 yielded an amplicon of 307 bp. For further screening, an *Afe1* restriction site was placed in the mutagenesis primer which digested the 307 amplicon into 109 bp and 198 bp fragments when the PAM[T^*3*55^A] mutation was present in either WT-AP53 cells or in AP5/*ΔsrtA* cells.

Similarly, to detect the *pam[T*^*355*^*Y]* mutation in WT-AP53 cells (AP53/*pam[T*^*355*^*Y*]) or AP53/*ΔsrtA* cells (AP53/*ΔsrtA/pam[T*^*355*^*Y*]), the same primers were used as for *pam[T*^*355*^*A]*, except that a new *NdeI* restriction site was present in the 307 bp amplicon. Digestion of the amplicon with *NdeI* provided 109 bp and 198 bp fragments when the *pam[T*^*355*^*Y]* mutation was present in either AP53 cells or AP53/*ΔsrtA* cells.

Also, PCR of the gDNAs using an external 5’-primer upstream of the *srtA* or *pam* gene with *cat-R*, and PCR of gDNAs with an external reverse primer downstream of the *srtA* or *pam* genes with *cat-F* provided correct amplicons in all cases, showing that *cat* and *pam genes* were appropriately targeted in each of the strains.

### Preparation of the CW, CM, and cytoplasm from AP53 cells

Overnight cultures of the different strains of AP53 cells were grown in Todd Hewitt medium supplemented with 1% yeast extract (THY) to mid-log phase (OD_600_ = 0.55-0.6) in an incubator set at 37° C/5% CO_2_. All strains were tested and confirmed for consistent and equal growth rates. The cultures (40 mL) were pelleted in a table-top centrifuge, and the supernates (200 µL) were taken for immunoassay with rabbit-anti-rPAM and concentrated 40x for Western analysis with rabbit-anti-PAM_short_.

The resulting pellets were washed 3X with PBS, then resuspended in 1 mL PBS-30% raffinose/20µg bacteriophage Ply-C (35), and incubated for 1 hr at room temperature. The resulting samples were then centrifuged for 10 min to provide the CW in the supernatants and the spheroplasts in the pellets. The CW fractions were removed and the spheroplasts were lysed by the addition of 1 mL sterile H_2_O, then vortexed to induce further lysis, and finally digested with 30 µL DNase I (2 units/µL) at 37° C for 30 min. Next, the samples were centrifuged at 165,000 x g for 3 hr at 4° C. The cytoplasm was removed as the supernatant and the CM pellet was then resuspended in 1 mL sterile H_2_O. The CW and CM samples were employed for Western blotting and fluorescent peptide cleavage assays.

### Cleavage of fluorescence-labeled peptides by recombinant (r)-sortases

Customized Dabcyl (4-((4-(dimethylamino)phenyl)azo)benzoic acid, succinimidyl ester)-LPSXG-(5-[(2-aminoethyl)amino]naphthalene-1-sulfonic acid) (Edans) peptides were provided by GenScript (Piscataway, NJ). Six peptide sequences were selected for study, *viz*., Dabcyl-LPS<u>T</u>GEAA-Edans, Dabcyl-LPS<u>A</u>GEAA-Edans, Dabcyl-LPS<u>E</u>GEAA-Edans, Dabcyl-LPS<u>K</u>GEAA-Edans, Dabcyl-LPS<u>Y</u>GEAA-Edans and Dabcyl-LPS<u>W</u>GEAA-Edans. The fluorescent peptides (0.2 mg) were dissolved in 1 mL 50 mM NaCl/50 mM Tris-HCl/0.2M NH_2_OH, pH 7.4. An aliquot of 100 µL of the peptide solution was transferred to replicate wells of a 96-well black fluorescence plate and 100 µL of 0.4 µM SrtA or SrtB was added to the individual wells. Wells in which buffer was substituted for sortases were used as controls. Fluorescence was measured at excitation and emission wavelengths of 350 nm and 495 nm, respectively, every 30 sec at 37° C. The change in fluorescence was calculated and the slope of the line by linear regression was determined using GraphPad Prism 9.

The same procedure was employed for cleavage of these peptides by the CM fractions of the AP53 isogenic cells.

### Binding of hPg to AP53 cells by Flow Cytometric Analysis (FCA)

Mid-log phase AP53 cells were grown as above. Bacteria cells were pelleted, washed twice in phosphate-buffered saline (PBS), and suspended in PBS/1% BSA blocking solution for 30 min. After this, ∼3 × 10^8^ cells were incubated with hPg (200 nM) for 60 min at 25° C. AP53 cells were washed twice with PBS and incubated for 30 min with monoclonal mouse-anti-hPg IgG (ERL, South Bend, IN). The cells were then washed twice with PBS and the AP53-bound hPg was detected by incubation with Alexa Fluor 488-donkey-anti-mouse IgG (Invitrogen) in the dark for 30 min. Finally, the cells were washed twice with PBS and fixed in PBS/1% paraformaldehyde. Fluorescence data were acquired, using the BD FACSAria III (BD Biosciences) by gating on fluorescence (FITC-A) and side-scatter, with scales set to logarithmic amplification. The cells in suspension were analyzed at a flow rate of 10 µL/min and 10,000 events per analysis. Histograms were analyzed using FCS Express Version 4 software (De Novo Software, Los Angeles, CA). The median fluorescence intensity (MFI), which measures the ability of each isogenic strain to bind hPg, was plotted against the respective strain using GraphPad Prism 9.

The binding of anti-PAM or anti-PAM_short_ to the isogenic AP53 cells was carried out in a similar way as described for hPg binding, except that the cells were incubated with rabbit-anti-rPAM polyclonal antibody for 30 min, then washed twice with PBS, and finally incubated with Alexa Fluor 488-chicken-anti-rabbit IgG (Invitrogen) in the dark for 30 mins. The succeeding steps were as above.

### Binding of anti-PAM to AP53 cells by whole cell ELISA

Mid-log phase AP53 cells were prepared as above. After 30 min incubation in PBS/1% BSA, ∼3×10^8^ cells were transferred to microcentrifuge tubes and incubated with rabbit-anti PAM_short_ for 30 min. The cells were centrifuged, washed twice in PBS, and followed by a 30 min incubation with goat-anti-rabbit IgG-HRP. Next, the cells were washed twice with PBS, resuspended in PBS, and the OD_600_ was adjusted to ∼0.16. Cell suspensions were diluted 5x and 50 µL of the suspensions were added in triplicate to individual wells of a 96-well microtiter plate. Color development was initiated by the addition of 100 µl HRP substrates, H_2_O_2_ and 3,3’,5,5’-tetramethylbenzidine (TMB). The reaction was allowed to proceed for 10 min at room temperature, after which it was terminated by the addition 50 µl 2N H_2_SO_4._ The A_450nm_ and OD_570nm_ were determined. The OD_570nm_ was subtracted from the A_450nm_ to correct for light scattering from bacterial cells. The data were analyzed by linear regression using GraphPad Prism software, version 9.0.

### Activation of hPg by AP53 whole cells

The procedure for the hPg continuous activation assay by SK2b has been described using mid-log phase AP53 cells. The bacteria were incubated in PBS/1% BSA for 30 min, and ∼2 ×10^8^ cells were added to individual wells of a 96-well microtiter plate. Blank wells contained PBS/1% BSA with no added cells. Next, hPg (400 nM) was added to all wells, followed by 15 min incubation. At this point, 100 µl of the chromogenic substrate, H-D-Val-L-Leu-L-Lys-p-nitroanilide (S2251) and SK2b in 10 mM Na-Hepes/150 mM NaCl, pH 7.4, were added to final concentrations of 0.25 mM and 5 nM, respectively. The final concentration of hPg was 200 nM. The hPm-catalyzed release of p-nitroaniline from S2251 was continuously monitored at 405 nm for up to 120 min. Initial velocities of activation were calculated as the slope of the linear region of a plot of A_405nm_ *vs* t^2^ (36). The data were analyzed using GraphPad Prism Version 9.

### Scanning Electron Microscopy (SEM) imaging of specific antibodies bound to AP53 GAS cells

Overnight cultures of isogenic AP53 GAS cells were grown at 37°/5% CO_2_ in THY medium until the mid-log phase was reached (OD = 0.55 - 0.60). All strains grew at approximately the same rates. The cultures (40 mL) were then pelleted using a routine tabletop centrifuge, washed with PBS, and then incubated for 1 hr with rabbit-anti-PAM, rabbit-anti-PAM_short_, or PBS. Subsequent to this step, the cells were then again pelleted and washed with PBS. The cell pellets were then placed in 2% glutaraldehyde/0.1M sodium cacodylate in PBS and 2 µL were spotted onto glass microscope slides coated with 0.01% poly-L-lysine in PBS for cross-linking to occur. The slides were then rinsed with PBS. Next, the slides were incubated for 1 hr in 1% OsO_4_ in PBS and then rinsed with PBS. The samples were then dehydrated stepwise using 50%, 70%, 80%, 95%, and 100% ethanol for 10 min. The ethanol was removed, and samples were submerged in liquid CO_2_ and allowed to reach the critical point to dry. The samples were then attached to SEM stubs and sputter coated with gold to 3 nm thickness. Samples were imaged at 80,000X using the FESEM-Magellan 400 (FEI).

## Data Availability

All data are supplied in this manuscript.

## References

1. Fischetti VA. 2016. M protein and other surface proteins on streptococci. In Ferretti JJ, Stevens DL, Fischetti VA (ed), Streptococcus pyogenes: Basic Biology to Clinical Manifestations, Oklahoma City (OK).

2. Pancholi V, Fischetti VA. 1992. A major surface protein on group A streptococci is a glyceraldehyde-3-phosphate-dehydrogenase with multiple binding activity. J Exp Med 176:415–426.

3. Pancholi V, Fischetti VA. 1998. alpha-enolase, a novel strong plasmin(ogen) binding protein on the surface of pathogenic streptococci. J Biol Chem 273:14503–14515.

4. Biagini M, Garibaldi M, Aprea S, Pezzicoli A, Doro F, Becherelli M, Taddei AR, Tani C, Tavarini S, Mora M, Teti G, D’Oro U, Nuti S, Soriani M, Margarit I, Rappuoli R, Grandi G, Norais N. 2015. The human pathogen Streptococcus pyogenes releases, lipoproteins as lipoprotein-rich membranevesicles. Mol Cell Proteomics 14:2138–2149.

5. Fischetti VA, Pancholi V, Schneewind O. 1990. Conservation of a hexapeptide sequence in the anchor region of surface proteins from gram-positive cocci. Mol Microbiol 4:1603–1605.

6. Denks K, Vogt A, Sachelaru I, Petriman NA, Kudva R, Koch HG. 2014. The Sec translocon mediated protein transport in prokaryotes and eukaryotes. Mol Membr Biol 31:58–84.

7. von Heijne G. 1989. The structure of signal peptides from bacterial lipoproteins. Protein Eng 2:531–534.

8. Carlsson F, Stålhammar-Carlemalm M, Flärdh K, Sandin C, Carlemalm E, Lindahl G. 2006. Signal sequence directs localized secretion of bacterial surface proteins. Nature 447:943–946.

9. Jeffery CJ. 2018. Protein moonlighting: what is it, and why is it importantã Philos Trans R Soc Lond B Biol Sci 373.

10. Hu P, Bian Z, Fan M, Huang M, Zhang P. 2008. Sec translocase and sortase A are colocalised in a locus in the cytoplasmic membrane of Streptococcus mutans. Arch Oral Biol 53:150–154.

11. Schneewind O, Missiakas DM. 2012. Protein secretion and surface display in Gram-positive bacteria. Phil Trans Royal Soc 367:1123–1139.

12. Mazmanian SK, Liu G, Ton-That H, Schneewind O. 1999. Staphylococcus aureus sortase, an enzyme that anchors surface proteins to the cell wall. Science 285:760–763.

13. Ton-That H, Liu G, Mazmanian SK, Faul KF, Schneewind O. 1999. Purification and characterization of sortase, the transpeptidase that cleaves surface proteins of Staphylococcus aureus at the LPXTG motif. Proc Natl Acad Sci USA 96:12424–12429.

14. Kumari P, Bowmik S, Paul SK, Biswas B, Banerjee SK, Murty US, Ravichandiran V, Mohan U. 2021. Sortase A: A chemoenzymatic approach for the labelling of cell surfaces. Biotechnol Bioeng doi:10.1002/bit.27935.

15. Russo BT, Ayinuola YA, Singh D, Carothers K, Fischetti VA, Flores-Mireles AL, Lee SW, Ploplis VA, Liang Z, Castellino FJ. 2020. The M-protein of Streptococcus pyogenes strain AP53 retains cell surface functional plasminogen binding after inactivation of the sortase A gene. J Bacteriol 202:e00096–e0020.

16. Jacobitz AW, Kattke MD, Wereszczynski J, Clubb RT. 2017. Sortase Transpeptidases: Structural Biology and Catalytic Mechanism. Adv Protein Chem Struct Biol 109:223–264.

17. Zhang Y, Liang Z, Glinton K, Ploplis V.A., Castellino FJ. 2013. Functional differences between Streptococcus pyogenes cluster 1 and cluster 2b streptokinases are determined by their β-domains. FEBS Lett 587:1304–1309.

18. Zhang Y, Liang Z, Hsieh HT, Ploplis VA, Castellino FJ. 2012. Characterization of streptokinases from group A Streptococci reveals a strong functional relationship that supports the coinheritance of plasminogen-binding M protein and cluster 2b streptokinase. J Biol Chem 287:42093–42103.

19. Grandi G. 2010. Bacterial surface proteins and vaccines. F1000 Biol Rep 2.

20. Steer AC, Law I, Matatolu L, Beall BW, Carapetis JR. 2009. Global emm type distribution of group A streptococci: systematic review and implications for vaccine development. Lancet Infect Dis 9:611–616.

21. Dale JB, Fischetti VA, Carapetis JR, Steer AC, Sow S, Kumar R, Mayosi BM, Rubin FA, Mulholland K, Hombach JM, Schodel F, Henao-Restrepo AM. 2013. Group A streptococcal vaccines: paving a path for accelerated development. Vaccine 31:B216–B222.

22. Fischetti VA. 2019. Vaccine Approaches To Protect against Group A Streptococcal Pharyngitis. Microbiol Spectr 7.

23. Bao YJ, Liang Z, Mayfield JA, Donahue DL, Carothers KE, Lee SW, Ploplis VA, Castellino FJ. 2016. Genomic characterization of a Pattern D Streptococcus pyogenes emm53 isolate reveals a genetic rationale for invasive skin tropicity. J Bacteriol 198:1712–1724.

24. Tokunaga M, Tokunaga H, Wu HC. 1982. Post-translational modification and processing of Escherichia coli prolipoprotein in vitro. Proc Natl Acad Sci USA 79:2255–2259.

25. Pancholi V, Chhatwal GS. 2003. Housekeeping enzymes as virulence factors for pathogens. Int J Med Microbiol 293:391–401.

26. Muller M, Koch HG, Beck K, Schafer U. 2001. Protein traffic in bacteria: multiple routes from the ribosome to and across the membrane. Prog Nucleic Acid Res Mol Biol 66:107–157.

27. Lee SG, Panchioli V, Fischetti VA. 2002. Characterization of a unique glycosylated anchor endopeptidase that cleaves the LPXTG sequence motif of cell surface proteins of Gram-positive bacteria. J Biol Chem 277:46912–46922.

28. Schneewind O, Missiakas D. 2019. Sortases, surface proteins, and their roles in Staphylococcus aureus disease and vaccine development. Microbiol Spectr 7.

29. Lee SG, Fischetti VA. 2006. Purification and characterization of LPXTGase from Staphylococcus aureus: the amino acid composition mirrors that found in the peptidoglycan. J Bacteriol 188:389–398.

30. Liang Z, Zhang Y, Agrahari G, Chandrahas V, Glinton K, Donahue DL, Balsara RD, Ploplis VA, Castellino FJ. 2013. A natural inactivating mutation in the CovS component of the CovRS regulatory operon in a pattern D Streptococcal pyogenes strain influences virulence-associated genes. J Biol Chem 288:6561–6573.

31. Raines RT, McCormick M, Van Oosbree TR, Mierendorf RC. 2000. The S.Tag fusion system for protein purification. Methods Enzymol 326:362–376.

32. Wang M, Prorok M, Castellino FJ. 2010. NMR backbone dynamics of VEK-30 bound to the human plasminogen kringle 2 domain. Biophys J 99:302–312.

33. Qiu C, Yuan Y, Liang Z, Lee SW, Ploplis VA, Castellino FJ. 2019. Variations in the secondary structures of PAM proteins influence their binding affinities to human plasminogen. J Struct Biol 206:193–203.

34. Nilsen SL, Castellino FJ. 1999. Expression of human plasminogen in Drosophila Schneider S2 cells. Protein Expr Purif 16:136–143.

35. Raz A, Fischetti VA. 2008. Sortase A localizes to distinct foci on the Streptococcus pyogenes membrane. Proc Natl Acad Sci USA 105:18549–18554.

36. Chibber BAK, Morris JP, Castellino FJ. 1985. Effects of human fibrinogen and its cleavage products on activation of human plasminogen by streptokinase. Biochemistry 24:3429–3434.

